# Diffusion approximations in population genetics and the rate of Muller’s ratchet

**DOI:** 10.1101/2021.11.25.469985

**Authors:** Camila Bräutigam, Matteo Smerlak

**Affiliations:** Max Planck Institute for Mathematics in the Sciences, Leipzig, Germany

## Abstract

The Wright-Fisher binomial model of allele frequency change is often approximated by a scaling limit in which selection, mutation and drift all decrease at the same 1/*N* rate. This construction restricts the applicability of the resulting “Wright-Fisher diffusion equation” to the weak selection, weak mutation regime of evolution. We argue that diffusion approximations of the Wright-Fisher model can be used more generally, for instance in cases where genetic drift is much weaker than selection. One important example of this regime is Muller’s ratchet phenomenon, whereby deleterious mutations slowly but irreversibly accumulate through rare stochastic fluctuations. Using a modified diffusion equation we derive improved analytical estimates for the mean click time of the ratchet.

## 1. INTRODUCTION

The Wright-Fisher (WF) model captures in simple mathematical form the combined effects of natural selection, mutation, and genetic drift, making it a pillar of population genetics. Unfortunately, the main quantities of interest, such as fixation times and fixation probabilities, cannot be computed analytically within this model—approximations are necessary. One established strategy is to build a diffusion process whose paths match those of the WF process over very long time scales. This *scaling limit* was first used by Fisher [1], Wright [2] and Kimura [3], and more rigorous expositions were given by Kolmogorov [4], Malécot [5] and Feller [6]; it now features prominently in textbooks under the name “Wright-Fisher diffusion equation” [7–10]. It has been said that “the standard diffusion approximation has permeated the field so thoroughly that it shapes the way in which workers think about the genetics of populations” [11].

Unfortunately, the classical WF diffusion equation has a rather narrow range of applicability: it can only account for evolutionary processes in which selection, mutation and genetic are all equally weak, scaling like the inverse population size 1/*N* (alternative approaches exist also for rescalings with 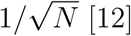). These conditions are neither biologically well motivated (selection coefficients and mutation rates do not covary with population size) nor universally applicable (viral populations, for instance, can have large sizes *O*(10^7^) [13]). Perhaps because the scaling limit of the WF model is often presented as *the* diffusion approximation of population genetics, diffusion methods are sometimes believed to be restricted to the weak selection-weak mutation regime of evolution. Outside this regime, the common practice is to either assume neutral evolution and effectively set all selection coefficients and mutation rates to zero, as in Kimura’s original work [3], or to use deterministic equations [14], *i*.*e*. to neglect genetic drift altogether.

Certain problems, however, require approximations that do better justice to the interplay between selection, mutation, and genetic drift. One example is *Muller’s ratchet* [15, 16], the irreversible accumulation of slightly deleterious mutations in finite asexual populations. Here, drift and the inflow of deleterious mutants drive the population to extinction against the force of natural selection. Muller’s ratchet has been invoked in the evolution of mitochondria [17], RNA viruses [18], Y chromosomes [19], ageing [20], cancer [21], and sex [22]. Obtaining analytical expressions for the mean click time of the ratchet has therefore been a longstanding goal for theory [19, 23–28].

At first sight, diffusion theory seems ideally suited for this problem: Muller’s ratchet is akin to a fixation problem for a two-type Wright-Fisher process, where one type represents unloaded alleles, and the other type comprises all other, deleterious mutations [19, 23, 26]. However, recent works have reinforced the belief that diffusion theory is inapplicable at finite selection strengths [27, 28]. An emerging consensus is that diffusion theory just does not apply to Muller’s ratchet, at least not in the slow click regime: “The fact that the extinction of the fittest class is due to […] a rare, large fluctuation and not […] a typical fluctuation prohibits simple diffusive treatments of the ratchet” [28]. We show that this belief is not justified. (As detailed in the Appendix, the approximation Eq. (23) in [28] can be in fact derived from the standard WF diffusion equation, Eq. (3).)

This paper has two objectives. First, we stress that diffusion methods in population genetics do *not* in fact require *s, u, v ∼* 1/*N* (selection coefficient *s*, mutation rates *u, v*) to capture the interplay between mutation and selection in large populations; an alternative diffusion equation, not based on taking a scaling limit, provides a more accurate approximation under the milder conditions *s, u, v ≪* 1 *≪ N*. The fact that diffusion theory can be used to derive more general results than the usual classical WF diffusion equation has been noted in the past (see e.g. [11] and references therein), but has not been sufficiently appreciated in our view. Second, we use the alternative approximation by diffusion to derive a new analytical formula for the mean fixation time of a deleterious allele, a central quantity in the context of Muller’s ratchet. We find that not only can diffusion theory be applied to this problem—it actually provides a better approximation than previous estimates in [27, 28] based on WKB expansions.

## II. TWO DIFFUSION APPROXIMATIONS

The haploid WF binomial model describes the evolution of the frequencies of two types *A* and *B*. For a population with size *N*, this process is the Markov chain on the set *{*0, 1/*N*, …, (*N −* 1)/*N*, 1*}* defined by

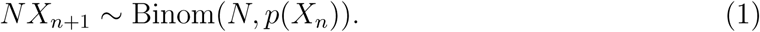

Here *X*_*n*_ denotes the frequency of *A* types at generation *n*, and *p*(*x*) represents the probability that an offspring is of type *A* given that the frequency of *A* in the parental population is *x*. The function *p*(*x*) depends on the specifics of mutation-selection dynamics and is parametrized by a selection coefficient and mutation rates. For instance, if *A* has (1 + *s*) times more offspring than *B*, and the probability of a mutation *A → B* (resp. *B → A*) is *u* (resp. *v*), we have

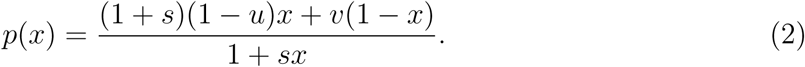

Other functional forms are more appropriate for viability selection, etc.

In the limit of large population sizes *N* → ∞, the standard diffusion approximation considers the WF model on the slow timescale *T* = *nN* and assumes that selection and mutation are comparable in strength to genetic drift, i.e. *Ns, Nu*, and *Nv* all have finite limits *σ, µ, ν* (respectively) as *N → ℞*. Under these conditions, it can be shown [7, 8] that the paths of the WF model on the slow timescale converge to those of a diffusion process with drift *σX*_*T*_ (1 *− X*_*T*_) *− µX*_*T*_ + *ν*(1 *− X*_*T*_)] and diffusion coefficient *X*_*T*_ (1 *− X*_*T*_). When brought back to the fast timescale *t* = *T/N*, this equation becomes

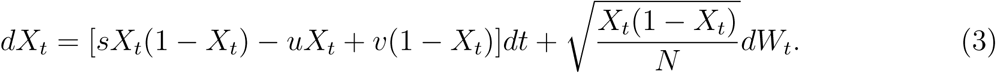

This is the “WF diffusion equation” from textbooks. (Here *dW*_*t*_ a standard Wiener process and the Ito convention is used.) By construction, this approximation requires that all evolutionary parameters scale as 1/*N* when *N* is taken to infinity, corresponding to the “weak selection-weak mutation” regime of evolution [7]. A rule of thumb due to Nei states that selection can be considered small if *Ns*^2^ *≪* 1 [27, 30].

Building a diffusion approximation of the WF process as a scaling limit implies that the deterministic term in (3) can only be linear in *s, u* or *v*: all higher order terms in *p*(*x*) in are suppressed in the large *N* limit. But according to (1) we have 𝔼 (*X*_*n*+1_ *− X*_*n*_|*X*_*n*_) = *p*(*X*_*n*_) *− X*_*n*_. It is therefore natural to consider as an alternative to (3) the diffusion equation

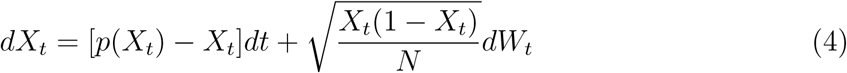

where the deterministic term is *not* linearized in *s, u, v*. We propose to use (4) to approximate of (1) whenever any of *s, u, v* in *p*(*x*) are small compared to one but large compared to 1/*N*. When *s, u, v* are in fact very small, (4) reduces to the classical WF diffusion (3). Eq. (4) appears in e.g. [26, 31], but, to our knowledge, has never been used to derive analytical results valid beyond the weak selection-weak mutation limit. A mathematical discussion of neutral diffusions at large mutation rates can be found in [32, 33].

## III. STATIONARY DISTRIBUTIONS AND FIXATION TIMES

### Stationary distributions

Given a stochastic differential equation of the kind described above,

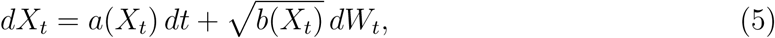

a stationary distribution *P*_eq_ exists as long as both boundaries, *x* = 0 and *x* = 1, are reflecting (*v* ≠ 0, *u* ≠ 0) and can be calculated as [34]

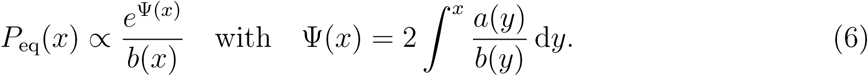

We evaluate this integral for the “old” (WF) diffusion equation (3) and the “new” diffusion (4) with sampling probability (2) to yield

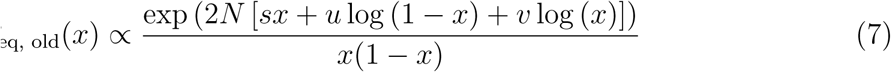

and

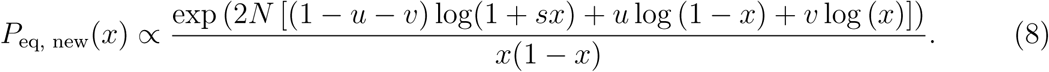

Fig. 1 compares these analytical predictions with empirical simulations of the Wright-Fisher model for a range of parameters *s, u, v*, both within and outside the weak selection-weak mutation regime defined by *s, u, v* = *𝒪* (1/*N*). Both the standard WF diffusion and new diffusion yield good approximations of the stationary distribution when *s <* 0.1. This supports the validity of the above expressions, for which we expect the predictions from the WF diffusion (in blue) and the new diffusion (in red) to be similar only while (1 *− u − v*) log 1 + *sx ≈ sx*. For stronger selection (*s ≥* 0.1), the WF diffusion severely overestimates the center of the distributions, which the new diffusion captures well. The new diffusion breaks down for *s* = 1 as expected.

**FIG. 1.**
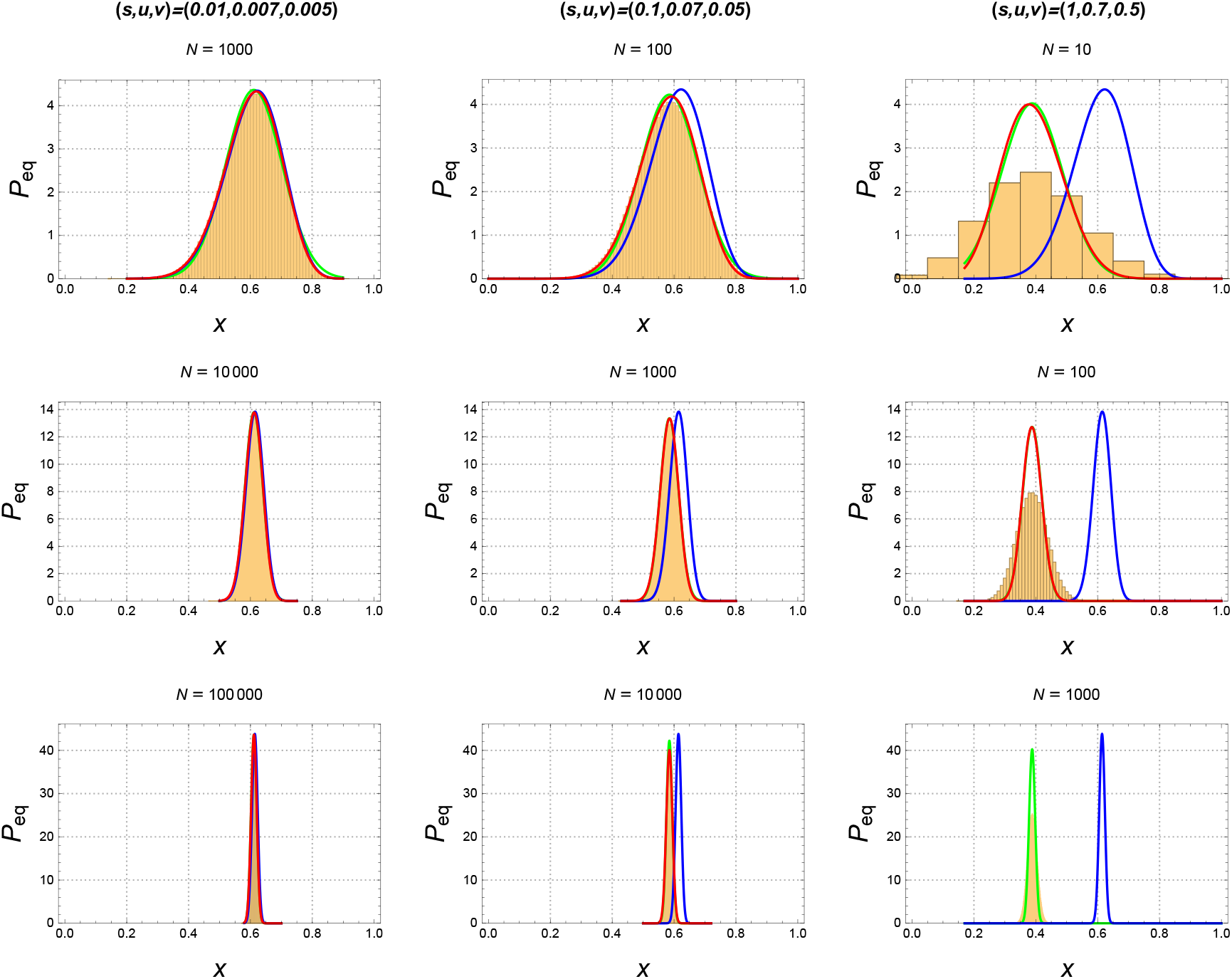
Stationary distributions from 10^6^ generations of the simulated WF process (Eq. (1), yellow histograms) compared to analytical predictions from the WF diffusion (Eq. (3), blue line), the diffusion by interpolation (Eq. (4), red line) and the prediction by Norman [29] (green line). In some cases the green and red line overlap completely so that only the red line remains visible. In the lower right plot the numerical integral of the new diffusion failed to converge. The parameters *s, u, v* as given in the subplots are kept constant when moving vertically downwards where population size *N* increases. Deviations of the WF diffusion from the underlying WF model become apparent for strong selection (*N ≫* 1/*s*^2^).

In Fig. 1 we additionally compare another approximation for the stationary distribution of the WF model, obtained by Norman in [29] (in green). Norman’s result predicts the stationary distribution to be Gaussian with mean *λ*, where *λ* is defined by the condition *a*(*λ*) = *p*(*λ*) *− x* = 0, and variance *σ*^2^ = *b*(*λ*)/(*N* |*a*^*′*^(*λ*)|), where *b*(*x*) = *p*(*x*)(1 *− p*(*x*)). The predicted stationary distributions are significantly better than the distributions derived from the standard WF diffusion approximation, but comparable in accuracy with the new diffusion. Note that we have considered that Norman’s derivation invoked a rescaling of time, so that the variance of the Gaussian distribution includes an additional factor of 2*N* compared to the expression in [29].

### Fixation probabilities

The classical fixation problem in population genetics considers the fate of a single selectively advantageous mutant (*s >* 0) subject to finite population size fluctuations (genetic drift) only. Mutation rates are zero (*u* = *v* = 0). Diffusive approximations can be used in that case to derive expressions for the probability of fixation at the boundaries *x* = 0 or *x* = 1.

The fixation probability using the new diffusive approximation, Eq. (4), is predicted to be

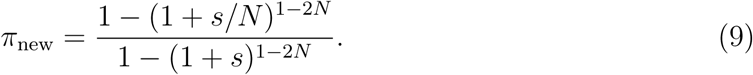

For comparison, the estimate from the standard WF diffusion (the Crow-Kimura formula [35]) reads

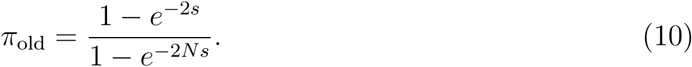

Again, these expressions give similar results while *s* is small so that *e*^*s*^ *≈* 1 + *s* holds.

We report a more complete discussion of the classical fixation problem, including fixation times and derivations in Appendix A, but note that the differences between predictions from the two diffusive schemes across varying *s* are only minor. This is due to the fact that fixation times depend largely on *s*, hence corrections with respect to *N* are small. When *s* is small, genetic drift is important but the two diffusions coincide. In contrast, for large *s* fixation will occur with high probability (and relatively fast), genetic drift does not play an important role.

The shortcomings of the standard WF diffusion approximation, compared to the new interpolation diffusion, become more apparent if the dynamics are not only determined by selection but also by counteracting back-mutations.

### Establishment times

To showcase this, we analyse the establishment of a beneficial mutant. Establishment occurs in a setting where a selectively advantageous mutant is additionally subjected to one-way reversion mutations (*u* ≠ 0, *v* = 0). A single advantageous mutant that appears in the population (*x*_0_ = 1/*N*) and survives genetic drift to reach a metastable state (*x*_*c*_ *>* 0, defined by *p*(*x*_*c*_) = *x*_*c*_) is said to have “established” [36]. In the long run the beneficial type will go extinct because *x* = 0 represents the only absorbing boundary. We are interested in probability and hitting time of establishment.

In Fig. 2 we compare the relative errors of mean establishment times *t*_est_ between WF simulations and predictions derived from the two diffusive approximations (Eq. (3) and Eq. (4)). Details of the calculations that yield the integral expression for the mean establishment times can be found in the Appendix B. The expressions need to be evaluated numerically. The comparisons from Fig. 2 confirm that for *s, u ≫* 1/*N* the competing forces of selection and mutation are better captured in the new diffusion scheme.

**FIG. 2.**
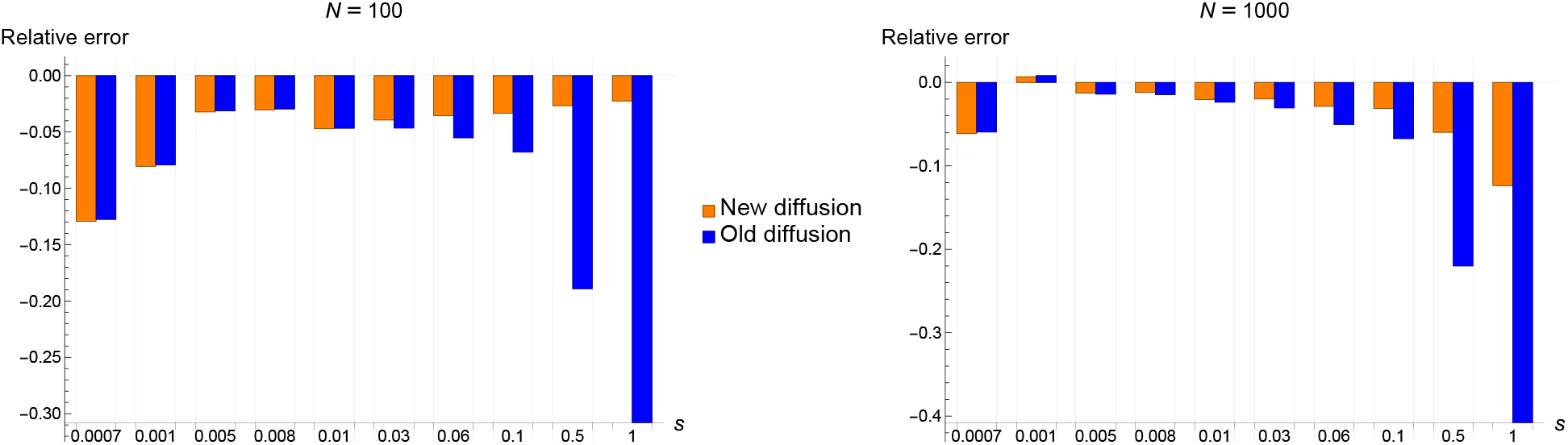
Relative errors in approximations of the establishment time of a rare mutant subject to reversions (here *u* = 5 *×* 10^*−*4^). The errors of the two analytical approximations, the standard WF diffusion (Eq. (3), in blue) and the alternative diffusion (Eq. (4), in orange) are computed relative to mean establishment times averaged across 10^6^ runs of stochastic WF simulations (Eq. (1)).

Nevertheless, the selection coefficients needed for a significant improvement compared to the standard WF diffusion are rather high. For most biological settings the WF diffusion would still perform sufficiently well. In the next section we find an example where the detailed interaction of selection, mutation and genetic drift matter for the dynamics of the population: the click time of Muller’s ratchet. This will highlight the improvements of the interpolation diffusion in biologically relevant parameter settings.

## IV. THE CLICK RATE OF MULLER’S RATCHET

The standard model of Muller’s ratchet (due to Haigh [37]) involves mutations occurring with rate *U* and with deleterious effect *S*, such that a mutant with *k* mutations has fitness (1 *− S*)^*k*^; the main quantity of interest in this context is the time to extinction of the un-mutated (*k* = 0) class. Waxman and Loewe [26] have shown that this problem can be simplified by lumping all *k*-mutants into one class with effective fitness (1 *− s*) where *s* = (1 *− e*^*−U*^)/(1 *− e*^*−U/S*^); the effective rate of mutation into this class is *u* = 1 *− e*^*−U*^. Within this framework, the click time of Muller’s ratchet is reduced to hitting time at *x* = 0 of the Wright-Fisher process defined by the sampling function

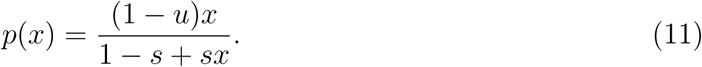

If mutations are not too frequent (*u < s*), the population will mostly equilibrate around a mutation-selection balance (*x*_eq_ = 1 *− u/s*), until a rare fluctuation drives the fit type to extinction against natural selection—Muller’s ratchet will click slowly. Note that the effective selection coefficient in the reduced model *s* can be larger than the biological coefficient *S*.

Using again classical expressions for hitting times of one-dimensional diffusions [7] together with Laplace’s approximation of integrals, we derive from Eq. (4) an analytical expression for the expected click time *T*_click_; see Appendix C for details. On a logarithmic scale, the resulting expression reads

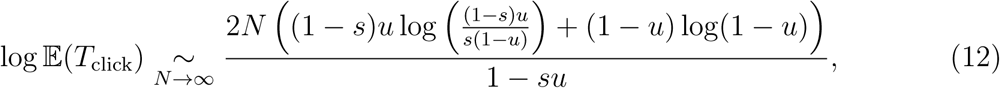

to be compared with the result using the standard WF diffusion Eq. (3)

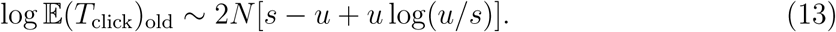

We plot the full analytical solution, Eq. (C5), in Fig. 3, together with simulations of the full Haigh model and the reduced stochastic WF model, as proposed by Waxman and Loewe [26]. We additionally compare in Fig. 3 the accuracy of our result with earlier heuristic diffusion approximations [38, 39] as well as more recent WKB expansions of Moran models [28]. We note that for the parameter ranges where the reduced model approximates the full model well, our analytical formula provides a very good fit. When the reduced model deviates, so does our analytical result. This simply suggests that the reduced model in the form as suggested in [26] cannot capture Haigh’s full model accurately across the whole parameter regime. Nevertheless, comparing to other analytical expressions derived for the full model, the prediction errors are small (the error of the reduced model is estimated to be within one order of magnitude [26]).

**FIG. 3.**
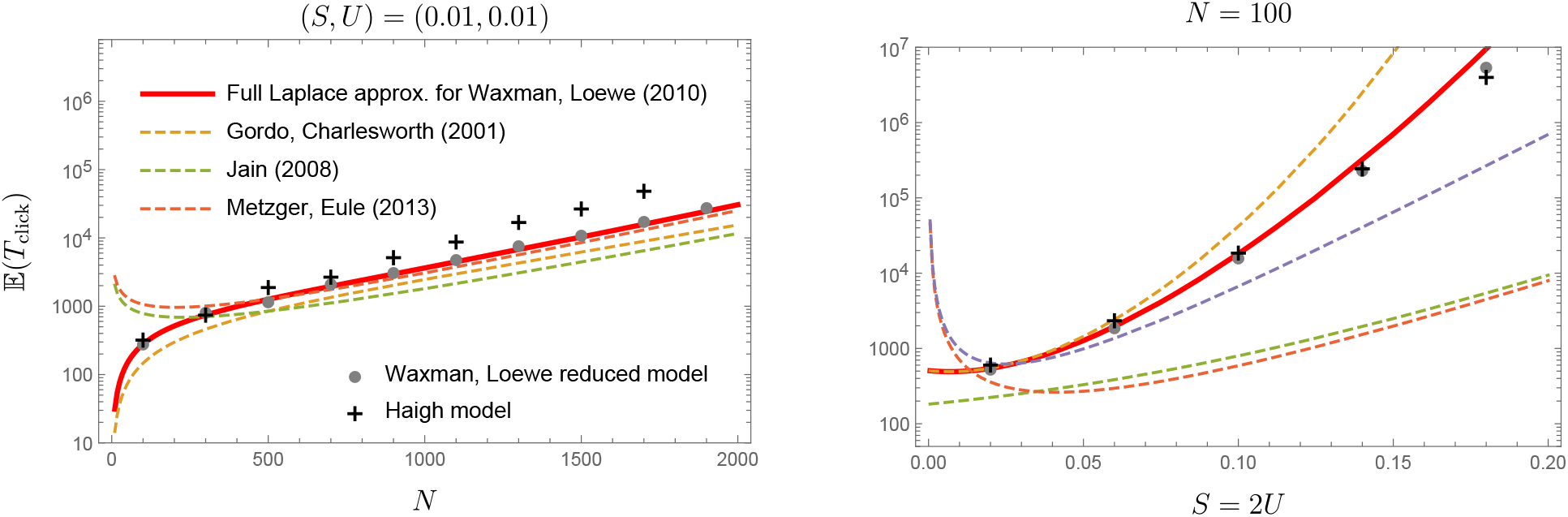
Accuracy of different analytical expressions for the mean time between clicks of the full model of Muller’s ratchet (black crosses), as a function of *N* for *S* = *U* = 0.01 (left) and as a function of *S* = 2*U* for *N* = 100 (right). Our result (Eq. (C5), red line) correctly approximates Wright-Fisher simulations of the two-type reduced model of Muller’s ratchet (mean click time over 10^3^ replicates, gray dots). Within one order of magnitude it also approximates click times of the full Haigh model.

Previous analytical approximations usually capture the rate of loss within narrow parameter ranges [38], with respect to a particular scaling (e.g. *s ∼* 1/*N*) [28, 39] or specifically for the fast clicking regime of Muller’s ratchet [23, 40]. We provide an analytical approximation that holds for a wider range of parameters, as presented in Fig. 3.

## V. CONCLUSION

We have argued that diffusion approximations of the Wright-Fisher model have broader range of applicability than usually appreciated from classical scaling arguments. In the context of Muller’s ratchet we found that Eq. (4) can be used to estimate the timescale of rare fixation events with good accuracy. This conclusion is in contrast with previously expressed concerns regarding the use of diffusion theory in the context of large deviations [41–44]: “The fact that the extinction of the fittest class is due to such a rare, large fluctuation and not the cause of a typical fluctuation prohibits simple diffusive treatments of the ratchet” [28]. Our results show, instead, that the analytical approximation of Muller’s ratchet within a diffusive framework improves previous results in the strong selection regime 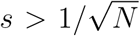. We expect that applications of diffusive approximations beyond the ones here presented might allow further progress in the analysis of evolutionary processes that are well captured by the discrete-time Wright-Fisher model, but not by its textbook diffusion approximation.

Since the mechanism of Muller’s ratchet was first suggested, experimental evidence for Muller’s ratchet as a fitness-decreasing effect on a population has accumulated in a variety of biological settings [45–47]. While our results show that this problem can be treated with standard diffusion tools, comparing our prediction with empirical data would require accurate estimate of evolutionary parameters, which is beyond the scope of this paper. Broadly, mutation rates per genome per generation have been estimated as large as *u*_*gen*_ *≈* 1 [48] for RNA viruses [49], hominids [50], inbred plant populations [51] or Drosophila [52]. Our model parameter *u* refers to the rate of deleterious mutations with specific fitness effect *s*, which are more difficult to extract experimentally. We chose to test our model on parameter ranges that have been discussed in the context of Muller’s ratchet [26, 27, 37, 38], as well as more recently in the attempt to quantify the specific threat of Muller’s ratchet in *C. Elegans* [53] and the Amazon molly [54].

We close with a semantic note. Because it does not depend on the mode (fecundity or viability) of selection, on generations being overlapping or non-overlapping, etc., the classical WF diffusion equation (3) is sometimes described as “universal”. What this means is simply that, in the special regime where *s, u, v* are all very small—as small as the inverse population size—, these differences are immaterial for the long-term dynamics of the population. But that is no longer true in regimes where selection and/or mutation are much stronger than genetic drift. In these regimes, diffusion theory does not break down; instead, other, non-universal diffusion approximations of the WF model can be used to obtain accurate results for fixation times and other quantities of interest.

## ACKNOWLEDGMENTS

We thank Maseim Kenmoe for help with Laplace integrals, Aleksander Klimek and Anton Zadorin for comments on the manuscript, and Sophie Pénisson for directing us to the mathematical references [32, 33]. We also thank two anonymous reviewers for helpful comments. Funding for this work was provided by the Alexander von Humboldt Foundation in the framework of the Sofja Kovalevskaja Award endowed by the German Federal Ministry of Education and Research.

## Appendix A: Fixation probability and fixation time

Eq. (4) from the main text describes the dynamics of the stochastic variable *X*_*t*_ in terms of a stochastic differential equation (SDE). Moving to an alternative dynamical description in terms of the probability density function *ρ*(*x, t*) of the stochastic variable *x* at time *t*, the corresponding drift-diffusion equation (also known as Fokker-Planck equation or Kolmogorov forward equation) takes the following form

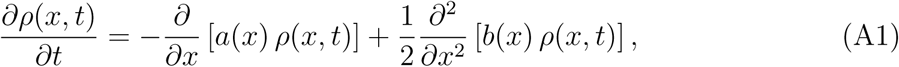

where in our setting *a*(*x*) = *p*(*x*) *− x* with *p*(*x*) as given in the main text (Eq. (2)), *b*(*x*) = *x*(1 *− x*)/*N*, with *x* being bound to the interval [0, 1] (since *x* describes a relative frequency).

For additional validation of our new model, we here compare predictions for the fixation probability *π* of a mutant with selection coefficient *s* in the Wright-Fisher model. The fixation problem considers the interplay of selection and genetic drift only. Mutation rates are assumed to be zero (*u* = *v* = 0).

Making use of Eq. (A1), the probability of fixation at *x* = 1, given an initial point *x* = *x*_0_, is approximated by

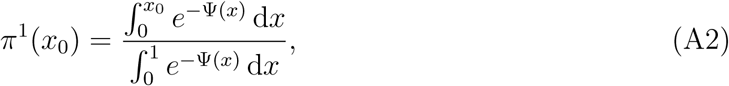

Where 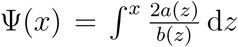. Similarly, the probability of fixation at *x* = 0, given an initial point *x* = *x*_0_, is approximated by

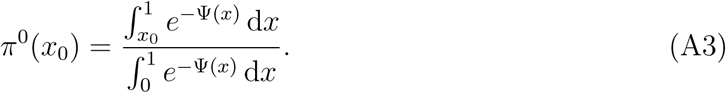

The classical fixation problem considers the probability of an initially single mutant *x*_0_ = 1/*N* to fix at *x* = 1.

Inserting *x*_0_ = 1/*N* into Eq. (A2) for the old and new diffusions diffusions gives the fixation probabilities given in the main text. In addition, Sella and Hirsch [55] suggest using

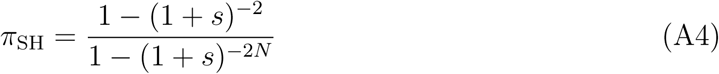

as a slightly more accurate estimate than the Crow-Kimura formula.

In Fig. 4 we compare both diffusive approximations (Eq. (3) and Eq. (4)), as well as the formula reported in [55] to empirical mean fixation probabilities from a total of 10^8^ (for *N* = 100) and 10^7^ (for *N* = 1000) Wright-Fisher simulations (Eq. (1)). Deviations between the old and new diffusive approximations only become apparent for strong selection *s*, as Eq. (9) and Eq. (10) are the same while *e*^*s*^ *≈* 1 + *s* holds. However, the improvements are only minor. For *s ≫* 0.1 we do not expect the new diffusion to be valid anymore and this also is supported by the decreased accuracy of prediction when selection coefficients become too large.

**FIG. 4.**
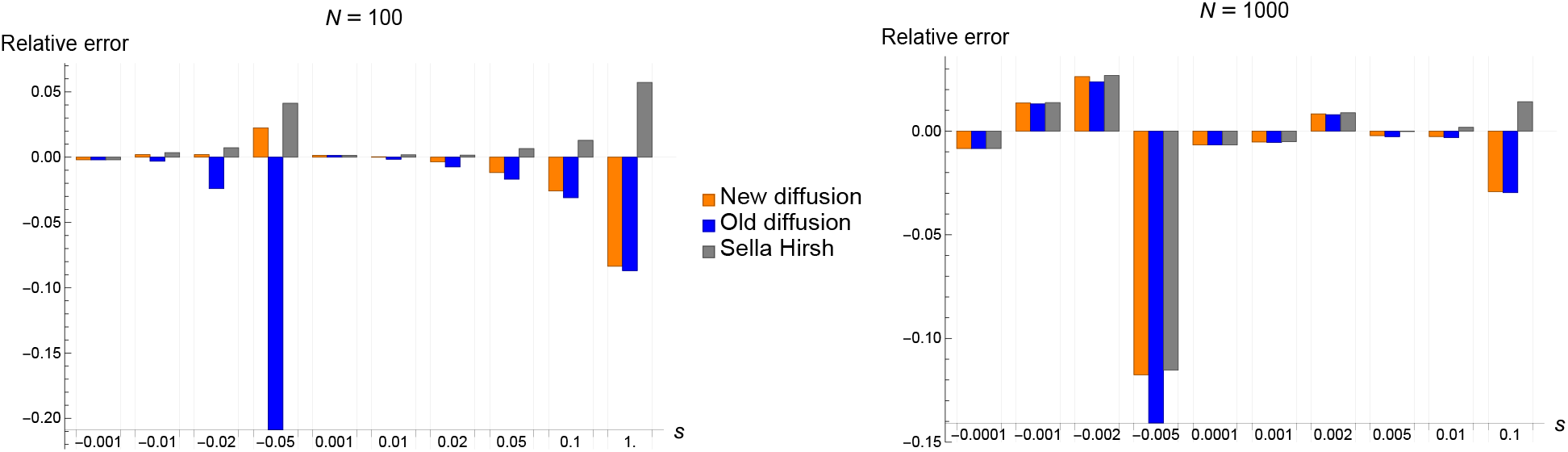
Errors in predicted fixation probabilities over varying selection coefficients *s* (*s <* 0: deleterious mutant, *s >* 0: advantageous mutant) and population sizes *N* = 100, 1000. Mutation rates are zero (*u* = *v* = 0). Approximations by Sella-Hirsh [55] (Eq. (A4), in grey), the WF diffusion (Eq. (10), in blue) [35], and from the new diffusion (Eq. (9), in orange) are compared to relative number of fixations in WF simulations.

Within the scope of diffusion theory we can also derive expressions for the mean fixation time. In the most general case, we are interested in the expected time for the stochastic variable *X*_*t*_ to hit the boundary at *x* = 0 or *x* = 1, given an initial position *X*_0_ = *x*_0_. This time 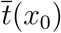 is described by the following second order differential equation [34]

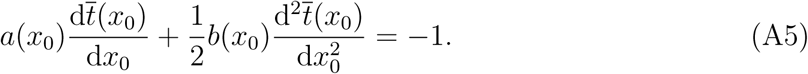

Solving Eq. (A5) subject to the appropriate boundary conditions 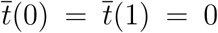 (two absorbing boundaries, since mutation rates are equal to zero), yields the mean first hitting time at *x* = 1, conditional on fixation at *x* = 1, with initial condition *x* = *x*_0_ [34],

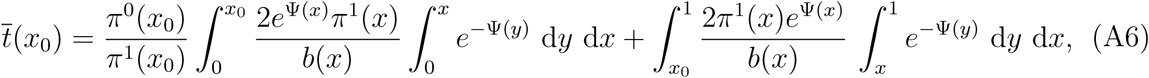

where as before 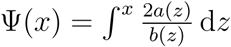, and *π*^1^(*x*) and *π*^0^(*x*) denote the fixation probabilities at *x* = 1 and *x* = 0, respectively.

The relative errors in predicting mean fixation times are compared in Fig. 5 between the WF diffusion (Eq. (3)) and the diffusive approximation by interpolation (Eq. (4)) relative to mean times of fixation from WF simulations for varying parameters *s, N*. The relative errors decrease substantially with increasing population size *N* in accordance with large *N* requirements for both diffusion approximations. While *s <* 0.1 the WF diffusion actually outperforms the new diffusion. We note, that this might be due to limited accuracy in the numerically evaluated integrals of Eq. (A6).

**FIG. 5.**
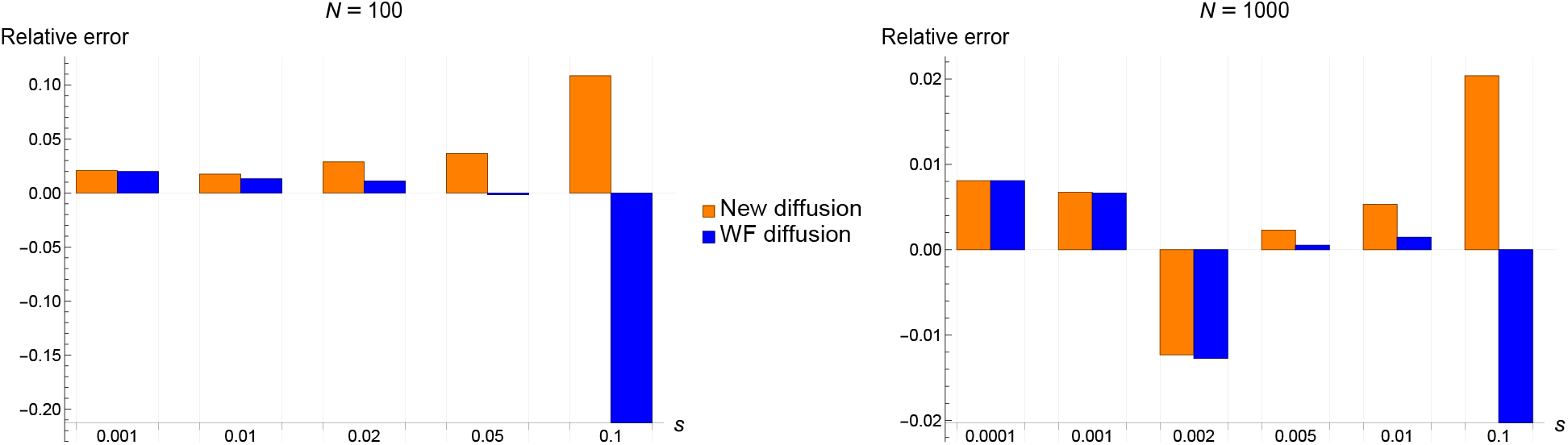
Relative errors in approximations of fixation times for varying selection coefficients *s* and population sizes *N* = 100, 1000 as calculated from Eq. (A6), for the WF diffusion (in blue) and the new diffusion (in orange). Errors are computed relative to simulated mean fixation times over 10^8^ (*N* = 100) and 10^7^ (*N* = 1000) runs of of WF simulations.

## Appendix B: Establishment times with one-way mutations

In Fig. 2 of the main text we compare analytical estimates for establishment times between the two diffusive approximations for the Wright-Fisher model with selection and one-way mutations (*s, u >* 0, *v* = 0). These expressions for the expected time to establishment can be derived in analogy to the above described fixation times. The probability of establishment, given an initial point *x*_0_, is given by

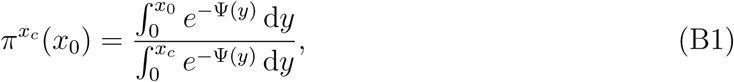

where as before 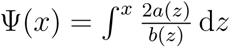 and *x*_*c*_ denotes the metastable state defined by *a*(*x*_c_) = 0. Note that the deterministic term *a*(*x*) now contains the parameter *u* ≠0 in contrast to the fixation problems discussed in the previous sections.

Assuming the initial frequency to be again a single mutant, 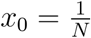, we can calculate the mean establishment time by evaluating the following integral numerically

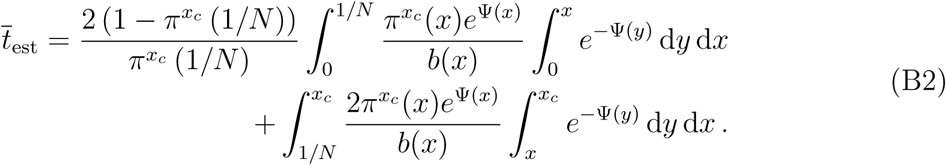

## Appendix C: Fixation of a deleterious mutant and Muller’s ratchet rate formula

The mean click time of Muller’s ratchet can be estimated by solving (A5) with sampling function (11) and boundary conditions

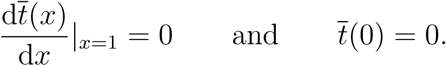

This gives

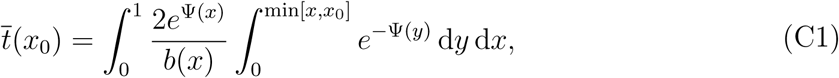

where 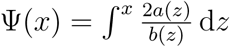 as before. Eq. (C1) cannot be solved analytically. Instead we note that the integral in the exponential, Ψ(*x*), is proportional to a large parameter, namely the population size *N*. This renders the integral suitable for a Laplace approximation in the so called low-noise regime *Ns ≫* 1. We therefore expand Ψ(*x*) in a Taylor series around the critical point *x*_*c*_ and truncate the expansion by assuming 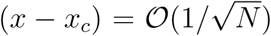. Hence we can write

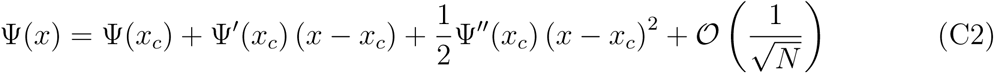

since 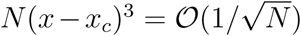. The exponential integral is reduced to a Gaussian integral on a bounded interval, which can be evaluated in terms of error functions. Generally speaking,

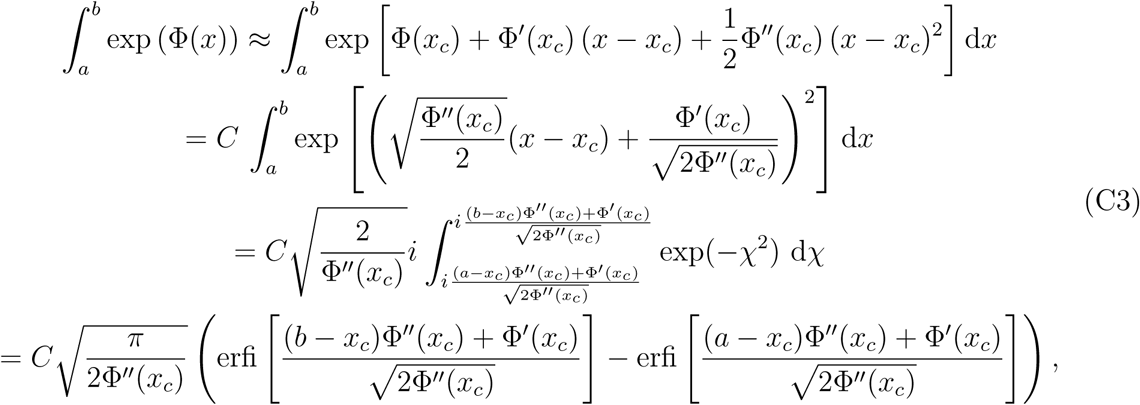

where 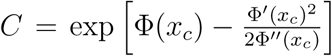 and erfi(*x*) refers to the imaginary error function. Considering in Eq. (C1) an additional expansion of the non-exponential function around the critical point 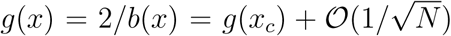, as well as an approximation of the limit min[*x, x*_0_] *≈* min[*x*_*c*_, *x*_0_] we notice that the previously nested integrals decouple. We can deduce the full approximation by applying the Laplace approximation (Eq. (C3)) to both integrals separately

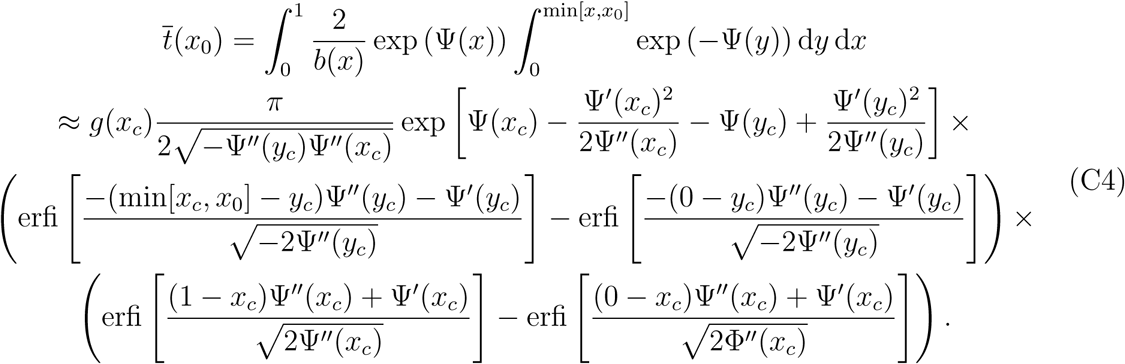

We assume the initial position to be precisely at the deterministic mutation-selection balance, i.e. where 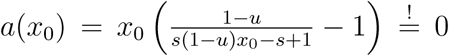, which means the initial point *x*_0_ and the critical point *x*_*c*_ of the potential Ψ(*x*) coincide, *x*_0_ = *x*_*c*_ = 1 *− u/s*. It is from this that the condition for the existence of an interior metastable state can be deduced, namely *u < s*. We arrive finally at the following expression for the expected click time in terms of our evolutionary parameters *N, s, u*:

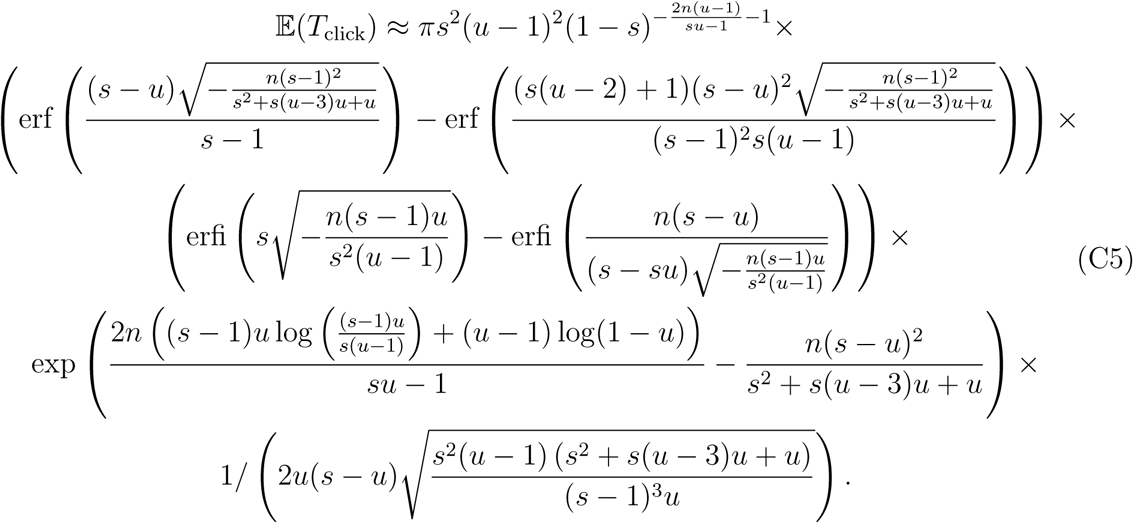

On a logarithmic scale, Eq. (C5) yields in the limit of large populations

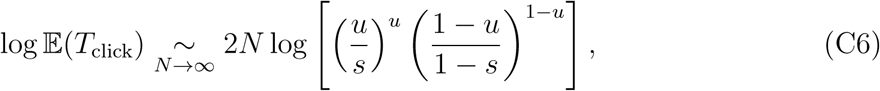

in accordance with Eq. (12) from the main text.

An analogous procedure can be used to derive an analytical expression for Muller’s ratchet rate from the standard WF diffusion. We consider the deterministic term of the WF diffusion (Eq. (3)) with *v* = 0, so that *a*(*x*) = *sx*(1 *− x*) *− ux*. This yields the following expected click time

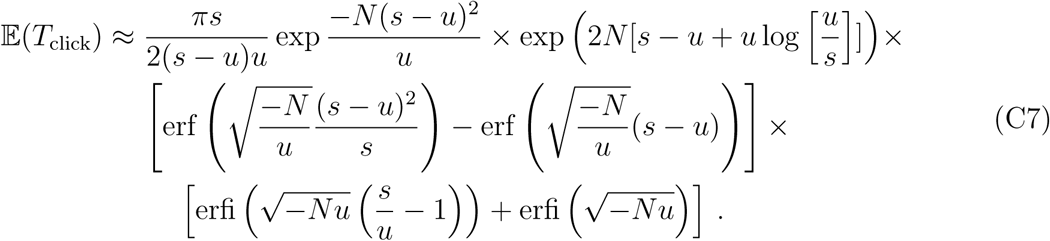

For *σ <* 1 *− u/s*, where *σ*^2^ = *u/*(2*Ns*^2^) denotes the variance of the Gaussian in the Laplace approximation, the expression from above can be reduced to

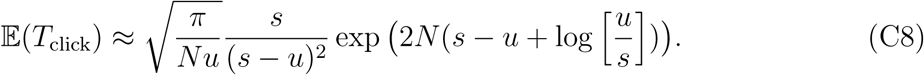

This is the same equation as derived in [28] through WKB approximations of the associated Moran process (after a rescaling of time and population size *t*^*′*^ *→ t/*2*N* and *N* ^*′*^ *→* 2*N*). The result in Eq. (C8) coincides on a logarithmic scale with Eq. (13), as described in the main text.

## Appendix D: Simulations

Stochastic simulations of the 2-type WF model were run in Wolfram Mathematica according to the dynamics of Eq. (1), while varying the evolutionary parameters *N, s, u, v* as described in the main text. Generated data points were usually averaged across 10^6^ repeats of the stochastic WF process or as specified otherwise. Only a constrained set of parameters were tested that did not exceed feasible computation times.

Simulations of a Haigh’s model of Muller’s ratchet [37], as well as the corresponding 2-type WF simulations of the reduced model [26], were implemented using the Julia programming language. For Haigh’s model, we initialised a monomorphic, mutation-free population and run the model until the mutation-free class was lost. As detailed in [37], the frequency of each class in the next generation is sampled from a multinomial distribution with sampling probability 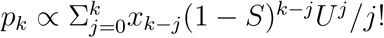.

The population in the reduced model evolves according to a WF model, as before, following the binomial sampling of Eq. (1), where *p*(*x*) = *x*(1 *− u*)/(1 *− s* + *s*(1 *− u*)). At the start of each simulation, the population is initialised at the mutation-selection equilibrium, as specified in the main text.

Our code is freely available at https://github.com/msmerlak/mullers-ratchet.

